# Harnessing multivariate, penalized regression methods for genomic prediction and QTL detection to cope with climate change affecting grapevine

**DOI:** 10.1101/2020.10.26.355420

**Authors:** Charlotte Brault, Agnès Doligez, Loïc le Cunff, Aude Coupel-Ledru, Thierry Simonneau, Julien Chiquet, Patrice This, Timothée Flutre

## Abstract

Viticulture has to cope with climate change and decrease pesticide inputs, while maintaining yield and wine quality. Breeding is a potential key to meet this challenge, and genomic prediction is a promising tool to accelerate breeding programs, multivariate methods being potentially more accurate than univariate ones. Moreover, some prediction methods also provide marker selection, thus allowing quantitative trait loci (QTLs) detection and allowing the identification of positional candidate genes. We applied several methods, interval mapping as well as univariate and multivariate penalized regression, in a bi-parental grapevine progeny, in order to compare their ability to predict genotypic values and detect QTLs. We used a new denser genetic map, simulated two traits under four QTL configurations, and re-analyzed 14 traits measured in semi-controlled conditions under different watering conditions. Using simulations, we recommend the penalized regression method Elastic Net (EN) as a default for genomic prediction, and controlling the marginal False Discovery Rate on EN selected markers to prioritize the QTLs. Indeed, penalized methods were more powerful than interval mapping for QTL detection across various genetic architectures. Multivariate prediction did not perform better than its univariate counterpart, despite strong genetic correlation between traits. Using experimental data, penalized regression methods proved as very efficient for intra-population prediction whatever the genetic architecture of the trait, with accuracies reaching 0.68. These methods applied on the denser map found new QTLs controlling traits linked to drought tolerance and provided relevant candidate genes. These methods can be applied to other traits and species.

## Introduction

Viticulture is facing two major challenges, coping with climate change and decreasing inputs such as pesticides, while maintaining yield and quality. This requires understanding the physiological processes and their genetic basis that determine adaptation to climate change, such as water use efficiency (Condon *et al.* 2004). Breeding schemes could then incorporate genotypes bearing genetic architecture favorable to high water use efficiency to be crossed with genotypes resistant to downy and powdery mildew (Vezzulli *et al.* 2019) and by selecting offspring combining favorable combinations. In crops, the widespread use of molecular markers through Marker Assisted Selection (MAS) or Genomic Prediction (GP) substantially accelerates the genetic gain compared to traditional phenotypic selection, by allowing early selection of promising genotypes, without phenotypic information (Heffner *et al.* 2009). This is of acute interest in fruiting perennial species because of the long juvenile period during which most traits of interest cannot be phenotyped. MAS and GP are now widely developed in many perennial species such as pear (Kumar *et al.* 2019), oil palm (Kwong *et al.* 2017; Cros *et al.* 2015), citrus (Gois *et al.* 2016), apple (Muranty *et al.* 2015) and coffea (Ferrão *et al.* 2019). In grapevine, QTL detection in biparental populations led up to the identification of major genes for traits with a simple genetic architecture such as resistance to downy and powdery mildew, berry color, seedlessness and Muscat flavor (Fischer *et al.* 2004; Welter *et al.* 2007; Fournier-Level *et al.* 2009; Emanuelli *et al.* 2010; Mejía *et al.* 2011; Schwander *et al.* 2012). Currently, most breeding effort in grapevine consists in improving disease resistance with MAS based on these results. However, genetic improvement is also needed for traits with more complex genetic determinism. Many minor QTLs have been found for the tolerance to abiotic stresses (Coupel-Ledru *et al.* 2014, 2016), yield components (Doligez *et al.* 2010, 2013) and fruit quality (Huang *et al.* 2012), as reviewed in Vezzulli *et al.* (2019). But MAS is not well suited for traits with many underlying minor QTLs (Bernardo 2008). Genomic prediction, which relies on high density genotyping is a promising tool for breeding for such complex traits, especially in perennial plants (Kumar *et al.* 2012). Nevertheless, in grapevine, GP has rarely been used yet, only once on experimental data (Viana *et al.* 2016a) and once on simulated data (Fodor *et al.* 2014). Thus, before applying GP to this species, it has to be empirically validated by thoroughly investigating the efficiency of different methods on traits with various genetic architectures.

Both QTL detection and genomic prediction rely on finding statistical associations between genotypic and phenotypic variation. So far, QTL detection in grapevine has been achieved mainly by using interval mapping (IM) methods in bi-parental populations, or more recently genome-wide association studies (GWAS) in diversity panels (see Vezzulli *et al.* (2019) for a comprehensive review of QTL detection studies in grapevine). However, most quantitative traits are explained by many minor QTLs which are hardly detected neither by interval mapping methods nor GWAS where each QTL has to individually overcome a significance threshold. In contrast, GP methods, by focusing on prediction, are less restrictive on the number of useful markers, sometimes resulting in all markers being retained as predictive with a non-zero effect. That is why GP methods are more efficient to predict genotypic values (Goddard and Hayes 2007) and therefore have become more and more popular with breeders (Heffner *et al.* 2010; Crossa 2017; Kumar *et al.* 2020).

Widely used methods for GP are based on penalized regression (Hastie *et al.* 2009), notably RR (Ridge Regression, equivalent to Genomic BLUP, GBLUP, Habier *et al.* (2007)) and the LASSO (Least Absolute Shrinkage and Selection Operator). Bayesian approaches are also commonly used in GP (e.g., de los Campos *et al.* (2013); Kemper *et al.* (2018)), see Desta and Ortiz (2014) for a classification of GP methods. However, Bayesian methods globally does not give better predictive ability than RR or LASSO, and they bear a heavy computational cost when fitted using Markov chains Monte-Carlo algorithms (Ferrão *et al.* 2019). Other methods based on non-parametric models (e.g., Support Vector Machine, Reproducing Kernel Hilbert Space, Random Forest) have been shown to yield lower predictive ability than parametric models (frequentist or Bayesian) when the genetic architecture of the trait was additive (Azodi *et al.* 2019).

Traits are often analyzed one by one in GP, using univariate methods. Nevertheless, breeders want to select the best genotypes which combine good performance for many traits. Analyzing several traits jointly in GP allows to take into account genetic correlation between traits (Henderson and Quaas 1976). Calus and Veerkamp (2011); Jia and Jannink (2012); Hayashi and Iwata (2013); Guo *et al.* (2014) compared univariate *vs* multivariate models’ performance. They found a slight advantage of multivariate analysis when heritability was low and data were missing. Predictive ability was particularly improved if the test set had been phenotyped for one trait while prediction was applied to another correlated trait (trait-assisted prediction) as in Thompson and Meyer (1986); Jia and Jannink (2012); Pszczola *et al.* (2013); Lado *et al.* (2018); Velazco *et al.* (2019); Liu *et al.* (2020). However, this breaks independence between the training and the test set, leading to an over-optimistic prediction accuracy (Runcie and Cheng 2019). Multivariate methods have also been proposed for QTL detection by Jiang and Zeng (1995); Korol *et al.* (1995); Meuwissen and Goddard (2004), notably for distinguishing between linkage and pleiotropy when a QTL is common to several traits. Some methods of multivariate penalized regression, such as in Chiquet *et al.* (2017), were designed to be useful for both QTL detection and prediction of genotypic value. Multivariate GP methods are expected to perform better if traits are genetically correlated but this remains worth testing with additional data. We also hypothesize that these methods will have higher power for QTL detection, by making a better use of information on the genetic architecture of several intertwined traits.

Methods designed for QTL detection are rarely used for genotypic value prediction. As they select only the largest QTLs, we hypothesize that these methods will provide an accurate prediction as long as the genetic architecture is simple, but yield poor prediction performance otherwise, as concluded in several studies (Heffner *et al.* 2011; Wang *et al.* 2014; Arruda *et al.* 2016). Conversely, some methods for GP like the LASSO and its extensions are also able to select markers with non-null effects, hence to perform QTL detection. Their accuracy in detecting QTLs has been partially investigated by Li and Sillanpää (2012) on a single trait in an inbred species and on simulated data and by Cho *et al.* (2010) on human data and binary trait, hence additional analyzes are needed.

This article aims to compare the ability of various methods to predict genotypic values and to detect QTLs in a bi-parental progeny of grapevine, by focusing on traits related to adaptation to climate change. We first complemented the available sparse SSR genetic map (Huang *et al.* 2012) by restriction-assisted DNA sequencing to construct a saturated SNP map. Then, we simulated phenotypic data using this map to compare several univariate and multivariate methods and assess the impact of simulation parameters. Finally, we reanalyzed the phenotypic data on water stress from Coupel-Ledru *et al.* (2014, 2016) obtained in semi-controlled conditions. The same genotyping data and methods as those applied to simulated data were compared, providing deeper insights into the genetic determinism of key traits underlying water use efficiency by finding new QTLs and candidate genes.

## Materials and Methods

### Plant Material

This study was based on a pseudo-F1 progeny of 188 offspring of *Vitis vinifera L.* from a reciprocal cross made in 1995 between cultivars Syrah and Grenache (Adam-Blondon *et al.* 2005).

### Genetic maps

#### SSR map

We used the genetic map with 153 multi-allelic SSR markers already published (Huang *et al.* 2012), constructed with the Kosambi mapping function (Doligez *et al.* 2013). In the following, we used the JoinMap version 3 format, according to which each marker genotype is encoded into one of the following segregation types: ab×cd, ef×eg, hk×hk, nn×np and lm×ll. Each of them comprises several allelic effects: e.g., for abxcd, the additive effects are a, b, c and d, and the dominance effects are ac, ad, bc and bd. Among the 153 SSRs, 50 were ab×cd, 58 ef×eg, 10 hk×hk, 16 lm×ll and 19 nn×np.

The physical positions of SSR markers absent from the latest URGI JBrowse (https://urgi.versailles.inra.fr/Species/Vitis/Genome-Browser) were retrieved by aligning their forward primer with BLAST (Altschul *et al.* 1990) on the PN40024 12X.v2 reference sequence (Canaguier *et al.* 2017) using default parameter values, except for the Expect threshold which was set to 1 or 10. Physical positions were still missing for six SSRs and one was uncertain (high Expect value).

#### SNP map

##### GBS markers

Further genotyping was done by sequencing after genomic reduction, using RAD-sequencing technology with *ApeKI* restriction enzyme (Elshire *et al.* 2011), as described in Flutre *et al.* (2020). Keygene N.V. owns patents and patent applications protecting its Sequence Based Genotyping technologies. This yielded a final number of 17,298 SNPs.

##### Consensus genetic map

The genetic map was built with Lep-MAP3 (Rastas 2017). The resulting map had 3,961 fully-informative markers (abxcd segregation) without missing data. These data were numerically recoded in biallelic doses (0,1,2) according to the initial biallelic segregation and phase (Table S1).

##### Design matrices

The resolution of multiple linear regressions described below requires a design matrix, which is built from the genotyping data. At a given marker, each genotype encoded in the JoinMap v3 format corresponded to several columns, yielding one predictor per allelic effect. From each genetic map (153 SSRs and 3,961 SNPs), we derived two design matrices, coded with 0, 1 and 2. The first one included only additive allelic effects (464 and 15,844, respectively). The second one included both additive and dominance allelic effects (996 and 31,688, respectively).

As mentioned before, we also recoded the 3,961 markers into additive gene dose (i.e., 0, 1 or 2), which yielded an additional design matrix with 3,961 predictors.

#### Simulation

Phenotype simulations were used to i) compare several methods for prediction accuracy, and ii) assess the efficacy of these methods to select the markers most strongly associated with trait variation.

Two traits, **y_1_** and **y_2_** were jointly simulated according to the following bivariate linear regression model: **Y** = **XB** + **E**, where **Y** is the *n × k* matrix of traits, **X** the *n × p* design matrix of allelic effects, **B** the *p × k* matrix of allelic effects, and **E** the *n × k* matrix of errors. For **X**, the 3,961 SNP markers mapped for the SxG progeny were used, encoded in additive and dominance effects. Therefore *n* = 188, *k* = 2, and *p* = 31,688. For **B**, allelic effects corresponding to *s* additive QTLs were drawn from a matrix-variate Normal distribution, **B** ~ *MV*(0, **I**, **V_B_**), with **I** being the *p* × *p* identity matrix and **V_B_** the *k* × *k* genetic variance-covariance matrix between traits such that 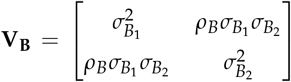 where *ρ_B_* is the genetic correlation among traits and 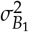 and 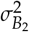 the genetic variances for both traits **y_1_** and **y_2_**. In the same way **E** ~ *MV*(0, **I**, **V_E_**), with the *k* × *k* error variance-covariance matrix 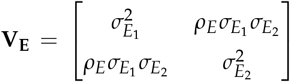 where *ρ_E_* is the residual error correlation among traits, and 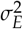 the error variance. We set *ρ_B_* to 0.8, 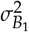 and 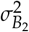 to 0.1, *ρ_E_* to 0 and narrow-sense heritability to 0.1, 0.2, 0.4 or 0.8 and 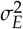 was deduced.

To explore different genetic architectures, we simulated *s* = 2 or *s* = 50 additive QTLs, located at *s* SNP markers, so that all corresponding additive allelic effects had non-zero values in **B**. Since all allelic effects were drawn from the same distribution, all QTLs had “major” or “minor” effects for *s* = 2 and *s* = 50, respectively. All dominant allelic effects were set to zero. Two QTL distributions across traits were also simulated. For the first one, called “same", all QTLs were at the same markers for both traits. For the second one, called “diff", the two traits had no QTL in common. Thus, there was genetic correlation among traits only for the “same” QTL distribution.

For each configuration (2 or 50 QTLs combined with “same” or “diff” distribution), the simulation procedure was replicated *t* = 10 times, each with a different seed for the pseudo-random number generator, resulting in different QTL positions and effects.

In a first simulation set, narrow-sense heritability was assumed equal for both traits and prediction was done with all methods. In a second set, we simulated two traits with different heritability values (0.1 and 0.5), for the “same” QTL distribution with *s* = 20 and *s* = 200 QTLs, with QTL effects drawn from a matrix-variate distribution with 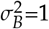 and *ρ_B_* = 0.5, in order to test the simulation parameters from Jia and Jannink (2012) with our genotyping data. For this second simulation set, prediction was done with a subset of methods only. Simulation parameters are summarized in Table 1.

**Table 1.** Parameter values in two sets of simulation of two traits in a bi-parental population

| Simulation parameter | Same heritability values | Different heritability values |
| --- | --- | --- |
| QTL number | 2-50 | 20-200 |
| Heritability value | 0.8/0.8 – 0.4/0.4 – 0.2/0.2 – 0.1/0.1 | 0.1/0.5 |
| Genetic variance | 0.1/0.1 | 1/1 |
| Genetic correlation | 0.8 | 0.5 |
| QTL distribution | Same-Diff | Same |

#### Experimental design, phenotyping and statistical analysis

Seven phenotypes related to drought tolerance had already been measured in two years on the Syrah x Grenache progeny (on 186 genotypes among the 188 existing) in semi-controlled conditions on the PhenoArch platform (https://www6.montpellier.inrae.fr/lepse_eng/M3P) in Montpellier, France, as detailed in Coupel-Ledru *et al.* (2014, 2016). Briefly, six replicates per genotype were used in 2012 (five in 2013). Three (in 2012) or two (in 2013) replicates were maintained under well-watered conditions (Well-Watered condition, WW), whereas the three other ones were submitted to a moderate water deficit (Water Deficit condition, WD). Specific transpiration, i.e. transpiration rate per leaf area unit, was measured during daytime (*TrS*) and night-time (*TrS*_*night*). Midday leaf water potential (*ψ_M_*, *PsiM*) was also measured and the difference between soil and leaf water potential (Δ*ψ*, *DeltaPsi*) was calculated. Soil-to-leaf hydraulic conductance on a leaf area basis (*KS*) was calculated as the ratio between *TrS* and *DeltaPsi*. Growth rate (*DeltaBiomass*) was estimated by image analysis. Transpiration efficiency (*TE*) was calculated over a period of 10 to 15 days as the ratio between growth and total water loss by transpiration during this period. These seven phenotypes were studied under each watering condition (WW and WD). We thus considered 14 traits in this study, a trait being defined as a phenotype x watering condition combination, and used the raw data available online (https://data.inrae.fr/privateurl.xhtml?token=383f6606-1c3c-4d90-8607-704cd53de068). For each trait, a linear mixed model was fitted with R/lme4 version 1.1-21 (Bates *et al.* 2014) using data from both years. First, a model with two random effects (genotype and genotype-year interaction) and nine fixed effects (year, replicate, coordinates in the platform within the greenhouse, coordinates in the controlled-environment chamber where *PsiM* and *TrS* were measured, operator for *PsiM* measurements, controlled-environment chamber and date of measurement) were fitted with maximum likelihood (ML). The best model among all sub-models was chosen using R/lmerTest version 3.1-2 (Kuznetsova *et al.* 2017) based on Fisher tests for fixed effects and likelihood ratio tests for random effects, with a *p*-value threshold of 0.05. This model was then fitted with restricted maximum likelihood (ReML) to obtain unbiased estimates of the variance components and empirical BLUPs (Best Linear Unbiased Predictions) of the genotypic values. The acceptability of underlying assumptions (homoscedasticity, normality, independence) was assessed visually by plotting residuals and BLUPs. Broad-sense heritability was computed according to Nanson (1970), dividing the residual variance by the mean number of trials (years) and replicates per trial. Its coefficient of variation was estimated by bootstrapping with R/lme4 and R/boot packages.

#### Interval Mapping methods

Two univariate interval mapping methods were compared, using R/qtl version 1.46-2 (Broman *et al.* 2003). For both, the probability of each genotypic class was first inferred at markers and every 0.1 cM between markers along the genetic map, using the R/qtl::calcgenoprob function.

##### Simple Interval Mapping

(SIM, Lander and Botstein (1989)) assumes that there is at most one QTL per chromosome. A LOD score was computed every 0.1 cM with R/qtl::scanone, then 1000 permutations were performed to determine the LOD threshold so that the family-wise (genome wide) error rate (FWER) was controlled at 5

##### Multiple Interval Mapping

(MIM, Kao *et al.* (1999)) allows the simultaneous detection of several QTLs. It was performed with R/qtl::stepwiseqtl, using a forward / backward selection of Haley-Knott regression model (Haley and Knott 1992), with a maximum number of QTLs set to 4 (or 10 for ROC curve construction, see below), replicated 10 times to overcome occasional instability issues. Only main effects were included (no pairwise QTL x QTL interaction). The LOD threshold was computed with permutations (1000 for QTL detection and 10 for cross-validation of GP, see below) to determine the main penalty with R/qtl::scantwo. QTL positions and effects were determined with R/qtl::refineqtl and R/qtl::fitqtl, respectively. For both methods, QTL positions were determined as those of LOD peaks above the threshold, with LOD-1 confidence intervals (Lander and Botstein 1989).

#### Penalized regression methods

Genomic prediction can be seen as a high-dimension regression problem with more allelic effects (in **B**) to estimate than observations (in **Y**), known as the “*n << p*” problem. The likelihood of such models must be regularized and various extensions, called penalized regression of the Ordinary Least Squares (OLS) algorithm were proposed. Such a penalization generally induces a bias in the estimation of allelic effects.

#### Univariate methods

##### Ridge Regression

(RR, Hoerl and Kennard (1970)) adds to the OLS a penalty on the effects using the *L*2 norm. As a result, all estimated allelic effects are shrunk towards zero, yet none is exactly zero. The amount of shrinkage is controlled by a regularization parameter (*λ*). We tuned it by cross-validation using the glmnet function of the R/glmnet package version 3.0-2 (Friedman *et al.* 2010) with default parameters except family = “gaussian” and *α* = 0, keeping the *λ* value that minimizes the Mean Square Error (MSE). Note that effects associated to correlated predictors are averaged so that they are close to identical, for a high level of regularization.

##### The Least Absolute Shrinkage and Selection Operator

(LASSO, Tibshirani (1996)) adds to the OLS a penalty on the effects using the *L*1 norm, causing some allelic effects to be exactly zero, while others are shrunk towards zero. Hence LASSO performs predictor selection, i.e., provides a sparse solution of predictors included in the best model, in addition to estimating their allelic effect. The LASSO regularization parameter (*λ*) was tuned by cross-validation with cv.glmnet (family = “gaussian”, *α* = 1). In the case of *n < p*, LASSO selects at most *n* predictors.

##### Extreme Gradient Boosting

Mason *et al.* (1999) is a machine learning method. We first applied the LASSO for reduction dimension and then Extreme Gradient Boosting to better estimate marker effect, based on the LASSO marker selection. Hence, we called that method LASSO.GB. As the LASSO estimation of allelic effect is biased, LASSO.GB could provide a better estimation, as well as the estimation of non-linear effects. Briefly, the Gradient Boosting iteratively updates the estimation of weak predictors, in order to reduce the loss. This method requires an optimization of many parameters associated with a loss function (MSE). This optimization has been done with train function from R/caret version 6.0-86 (Kuhn 2008) using the “xgbTree” method. As the optimization of numerous parameters was computationally heavy, we fixed some of them (nrounds = max_depth = 2, colsample_bytree = 0.7, gamma = 0, min_child_weight = 1 and subsample = 0.5), while testing a grid of varying parameters (nrounds = 25, 50, 100, 150; eta = 0.07, 0.1, 0.2).

##### The Elastic Net

(EN, Zou and Hastie (2005)) adds to the OLS both *L*1 and *L*2 penalties, the balance between them being controlled by a parameter (*α*). Both *α* and *λ* were tuned by a nested cross-validation: 20 values of *α* were tested between 0 and 1 and, for each of them, we used cv.glmnet function (from R/glmnet package) to choose between 500 values of *λ*. The parameter pair minimizing the MSE was kept. EN performs predictor selection but is less sparse than LASSO.

Note that RR, LASSO and EN all assume a common variance for all allelic effects.

#### Multivariate methods

##### The multi-task group-LASSO

(MTV_LASSO, Hastie and Qian (2016)) is a multivariate extension of the LASSO, *λ* parameter was tuned using glmnet (family = “mgaussian”, *α* = 1). It assumes that each predictor variable has either a zero or a non-zero effect across all traits, allowing non-zero effects to have different values among traits. MTV_RR is the multivariate extension of RR, also tuned with glmnet (family = “mgaussian”, *α* = 0). Similarly, MTV_EN is the multivariate extension of EN. The implementation of these three methods is identical.

##### The multivariate structured penalized regression

(called SPRING in Chiquet *et al.* (2017)) applies a *L*1 *penalty* (*λ*_1_ parameter) for controlling sparsity (like LASSO) and a smooth *L*2 *penalty* (*λ*_2_ parameter) for controlling the amount of structure among predictor variables to add in the model, i.e., the correlation between markers according to their position on the genetic map. Both parameters were tuned by cross-validation using cv.spring function (from R/spring package, version 0.1-0). Unlike multitask group-LASSO, SPRING selects specific predictors for each trait, i.e., a selected predictor can have a non-zero effect for a subset of the traits. SPRING allows the distinction between the direct effects of predictors on a trait and their indirect effects by using conditional Gaussian graphical modeling. These effects are due to covariance of the noise such as environmental effects affecting several traits simultaneously. This distinction results in two kinds of estimated allelic effects: the direct ones, re-estimated with OLS, which are best suited for QTL detection (we called the corresponding prediction method **spring.dir.ols**) and the regression ones, which involve both direct and indirect effects and are best suited for prediction (**spring.reg** method).

#### Robust extension for marker selection

To enhance the reliability of marker selection by penalized methods, we used two approaches: Stability Selection (Meinshausen and Buhlmann 2009) and marginal False Discovery Rate (Breheny 2019), which both aim at controlling the number of false positive QTLs. We did not use these methods for genomic prediction, as they are not designed for this purpose.

##### Stability selection

(SS) is a method which controls the FWER, computes the empirical selection probability of each predictor by applying a high-dimensional variable selection procedure, e.g., LASSO, to a different subset of half the observations for each *λ* value from a given set, and then keeps only predictors with a selection probability above a user-chosen threshold. Stability selection is implemented in R/stabs package version 0.6-3 (Hofner and Hothorn 2017) and can also be adapted to a multivariate framework. For QTL detection on experimental data, the probability threshold we applied was 0.6 for LASSO.SS and 0.7 for MTV_LASSO.SS.

##### Marginal False Discovery Rate

(mFDR) allows to choose a more conservative value of *λ* for LASSO and EN with the R/nvcreg package version 3.12.0 (Breheny 2019). For QTL detection on experimental data, we set mFDR to 10% for LASSO.mFDR and EN.mFDR. This approach is not adapted to a multivariate framework.

#### Evaluation and comparison of methods

All methods were compared on two aspects: their ability to predict genotypic values, and their ability to select relevant markers, i.e., to detect QTLs. To assess the prediction of genotypic values on simulated data, we used the Pearson’s correlation coefficient between the predicted genotypic values and the simulated ones (prediction accuracy). On experimental data, we used the same criterion, but the true genotypic values being unknown, we used their empirical BLUPs instead (predictive ability).

For QTL detection on simulated data, the methods were compared using criteria of binary classification based on the numbers of true positives and false negatives. On experimental data, because true QTLs are unknown, no such comparison could be performed; instead, we compared the outcome of the different methods and QTLs were deemed reliable when found by several methods.

#### Genomic prediction

A nested cross-validation (CV) was applied to assess prediction by the various methods.

- An outer **k1***fold* CV was performed to estimate the performance metrics, with an inner **k2** − *f old* CV applied to the training set of each outer fold to find the optimal tuning parameters for the method under study (Figure S2). Both **k1** and **k2** were set to 5 (see Arlot and Lerasle (2016). The folds of the outer CV were kept constant among traits and methods.
- For interval mapping methods, the optimal tuning parameter was the LOD threshold obtained from permutations, and the effects for the four additive genotypic classes (ac, ad, bc and bd) were estimated by fitting a multiple linear regression model with genotype probabilities at all QTL peak positions as predictors, using R/stat::lm. For penalized regression methods, parameters were optimized with specific functions such as cv.glmnet and cv.spring.
- As performance metrics, we used mainly the Pearson’s correlation (corP) but we also calculated the root mean square predicted error (RMSPE), the Spearman correlation (corS), the model efficiency (Mayer and Butler 1993) and test statistics on bias and slope from the linear regression of observations on predictions (Piñeiro *et al.* 2008).

For experimental data, the whole nested cross-validation process was repeated 10 times (*r*=10), whereas for simulated data it was performed only once, but on 10 different simulation replicates (*r*=1 and *t*=10). The 14 traits were analyzed jointly for MTV_RR, MTV_LASSO and MTV_EN. But for SPRING, since analyzing all traits together was computationally too heavy, we split traits into three groups by hierarchical clustering (Figure S3) performed with R/hclust applied to genotypic BLUPs. All traits within each group were analyzed together.

For simulated data with the same heritability values for both traits, performance results were averaged not only over simulation replicates and partitions of outer CV, but also over traits, because both traits were equivalent in terms of simulation parameters. For simulated data with different heritability values, performance results were averaged only over simulation replicates and partitions of outer CV. For experimental data, performance results were averaged over partitions of outer CV and outer CV replicates.

Moreover, in terms of design matrices, for experimental data, we compared several ones based on the mean predictive ability of eight methods across the 14 traits of experimental data. For IM methods, only SSR and SNP maps coded in JoinMap format were compared. We showed that for most methods, the SNP genotypes recoded into gene doses gave the best predictive ability (Figure S4), tied with other SNP design matrices. For computational reasons, we hence chose to use this one for method comparison. For simulated data, as QTLs correspond to SNP markers, we only used the SNP map as the design matrix, coded in gene doses for penalized methods and in JoinMap format for IM methods.

#### QTL detection

##### Simulated data

The quality of a predictor selection method is usually assessed with the relationship between statistical power (i.e. the True Positive Rate, TPR) and type I error rate (i.e. the False Positive Rate, FPR). To compare methods, we thus used the ROC (receiver operating characteristic) curve (Swets *et al.* 1979), which is the plot of TPR as a function of FPR over a range of parameters (Table 2), and the pAUC (partial Area Under the Curve; McClish (1989); Dodd and Pepe (2003)). Any marker selected at +/− 2 cM of a simulated QTL was counted as a True Positive.

**Table 2.** Parameter ranges for ROC curve computation for comparing predictor selection performance of different methods.

| Method | SIM / MIM | LASSO / MTV_LASSO | Stability Selection | SPRING | EN | mFDR |
| --- | --- | --- | --- | --- | --- | --- |
| Parameter name | LOD | $\lambda$ | probability threshold | $\lambda_1$ | $\lambda$ | mFDR |
| Lowest constraint | 0 | 10e-5 | 0.5 | 10e-8 | 10e-4 | 0.3 |
| Highest constraint | 14 | 0.25 | 0.9 | 0.25 | 8 | 0 |

For methods with two tuning parameters, one parameter was kept constant (*α* at 0.7 for EN and EN.mFDR, and *λ*_2_ at 10e-8 for SPRING). We tested several values of *α* for EN but it did not change much the results (not shown). For MIM, the maximum number of QTLs that can be integrated into the model was set to 10.

##### Experimental data

Comparison between methods was based on the number of detected QTLs, the magnitude of their effects and the percentage of variance globally explained by all detected QTLs.

For MTV_LASSO and SPRING, we split traits into three groups as described above, for computational reasons (for SPRING) and to test whether such splitting gave more reliable QTLs (for MTV_LASSO). The parameters of penalized methods were tuned by cross-validation, with MSE as the cost function. We compared predictor selection between methods in terms of the number of common selected markers per trait, i.e. the intersection between markers selected by penalized methods (focusing on LASSO and EN) and markers inside confidence intervals found by interval mapping methods (focusing on MIM). Then all markers in high LD with those selected were considered as selected too. The threshold was defined as the 95% quantile of LD value distribution, for all pairs of markers belonging to the same chromosome (Figure S5), which gave a LD threshold of 0.84.

We deemed selected markers as highly reliable if they were either i) selected by at least five methods, whatever the methods, ii) or selected by both EN.mFDR and MIM (see Results). Then, we defined a highly reliable QTL as the interval of +/− 3 cM around each highly reliable marker (Price 2006; Viana *et al.* 2016b), as predicted by loess fitting of genetic positions to physical positions. When several markers were selected inside the 6 cM interval, the QTL interval was extended accordingly. The genetic positions of this interval were then converted into physical positions, by fitting a polynomial local regression (loess). QTLs overlapping for several traits on the SNP map were merged into a single QTL, by physical intervals’ union. We determined QTLs overlapping between SSR and SNP genetic maps based on physical positions.

##### Candidate genes exploration

After merging the most highly reliable QTLs colocalized between traits, we proceeded to search for underlying candidate genes. We retrieved the list of genes overlapping the intervals of our QTLs from the reference Vitis genome 12X.v2 and the VCost.v3 annotation (Canaguier *et al.* 2017). We then used the correspondence between IGGP (International Grapevine Genome Program) and NCBI RefSeq gene model identifiers provided by URGI (https://urgi.versailles.inra.fr/Species/Vitis/Annotations) to get putative functions from NCBI, when available. For those genes with a putative function, we then refined the analysis to retrieve additional information about their function and expression. We searched UniProt (https://www.uniprot.org/) and TAIR (https://www.arabidopsis.org/) databases to get a complete description of the genes function, their name and the corresponding locus in Arabidopsis. In addition, we used the GREAT (GRape Expression Atlas) RNA-seq data analysis workflow (https://great.colmar.inrae.fr/app/GREAT), which gathers published expression data, to assess the level of expression of our candidate genes in grapevine leaves and shoots, the organs relevant for the traits considered in this study. RNA-seq data are normalized as detailed on the ‘User manual’ section of the GREAT platform: “from the raw read counts, the normalized counts (library size normalization) and the RPKM (gene size normalization) are calculated for each gene in each sample”. Data were retrieved with all filters set to “Select All” except for the organ considered that was restricted to ‘Leaves’ and ‘Shoot’.

#### Data availability and reproducibility

All software we used was free and open-source and most analyzes were done with R (R Core Team 2020), notably graphs were created using the ggplot2 package (Wickham 2016). All R scripts used for the analysis, i.e. genetic mapping, simulation, phenotypic analysis, prediction and QTL detection, are available in a first, online repository at https://data.inrae.fr/privateurl.xhtml?token=d7ef7492-a2a7-499d-82c0-baad1d14a8dd. Many of the custom functions we used are available in a package for reproducibility purposes, R/rutilstimflutre (Flutre 2019). Raw and transformed genotypic data, as well as the genetic map, are available in a second, online repository at https://data.inrae.fr/privateurl.xhtml?token=782ff6ff-d79c-4714-b0da-b85c5a4514a5.

## Results

### Genetic mapping

We constructed a saturated consensus genetic map with 3,961 SNP markers obtained by GBS. The SNP map covers 1,283 cM. It was essentially superimposed on the SSR map of 1,116 cM (Figure 1). Chromosome 17 had the largest contribution to this 15% difference in length, its length being 37.8 cM with SSRs and 63.7 cM with SNPs. Chromosomes 2, 3, 12, 13 and 15 were also longer on the SNP map. The average distance between markers was 0.34 cM for the SNP map (respectively 9.0 cM for the SSR map) and the maximum distance was 12.0 cM (respectively 29.4 cM for the SSR map). At most places along the genome, genetic order was consistent with physical order.

**Figure 1.**
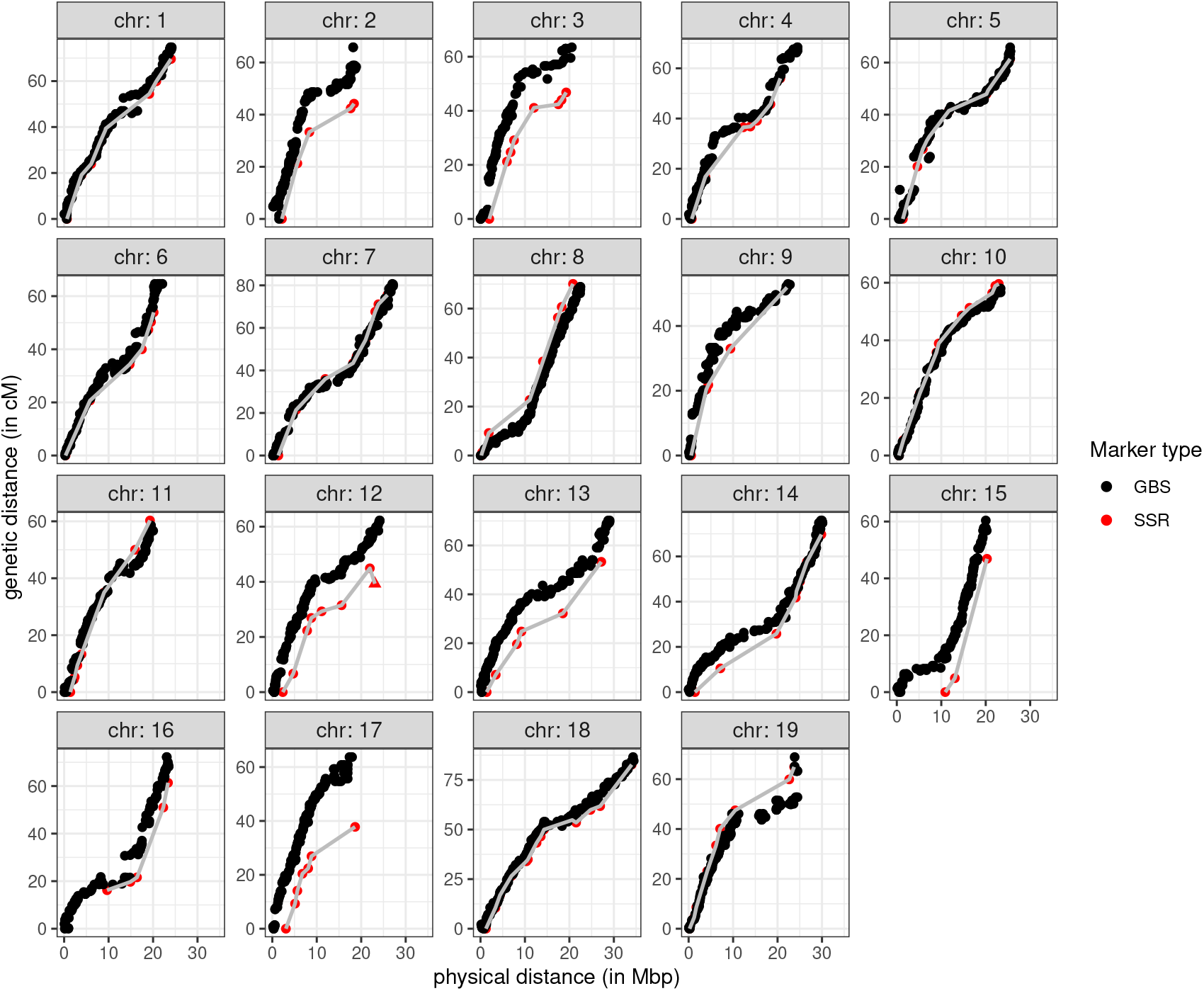
Comparison of SSR and SNP consensus genetic maps of a pseudo-F1 *V. vinifera* population, obtained by plotting genetic positions as a function of physical positions for each chromosome. The position of the SSR marker indicated by a triangle on chromosome 12 was uncertain.

### Comparison of methods with simulated data

#### Prediction: cross-validation results

##### Traits with the same heritability value

Methods were compared for prediction accuracy by applying cross-validation on simulated data with four different configurations and four heritability values.

Mean Pearson’s correlation coefficient varied from 0.16 to 0.98, with a strong effect of heritability on prediction accuracy in all configurations, for the seven main methods (Figure 2). As expected, MIM performed very well in the “major” configurations across all heritability values but yielded the least accurate prediction in the “minor” ones. On the opposite, RR performed very well in the “minor” configurations, but yielded the least accurate prediction in the “major” ones. EN prediction performance was always intermediate between those of RR and LASSO. QTL distribution among traits - “same” (for QTLs at the same positions) or “diff” (for QTLs at different positions) - had very little effect on prediction accuracy. Moreover, we did not observe any superiority of multivariate methods over univariate ones, despite the strong genetic correlation simulated between traits (*ρ_B_*=0.8) and no correlation between errors.

**Figure 2.**
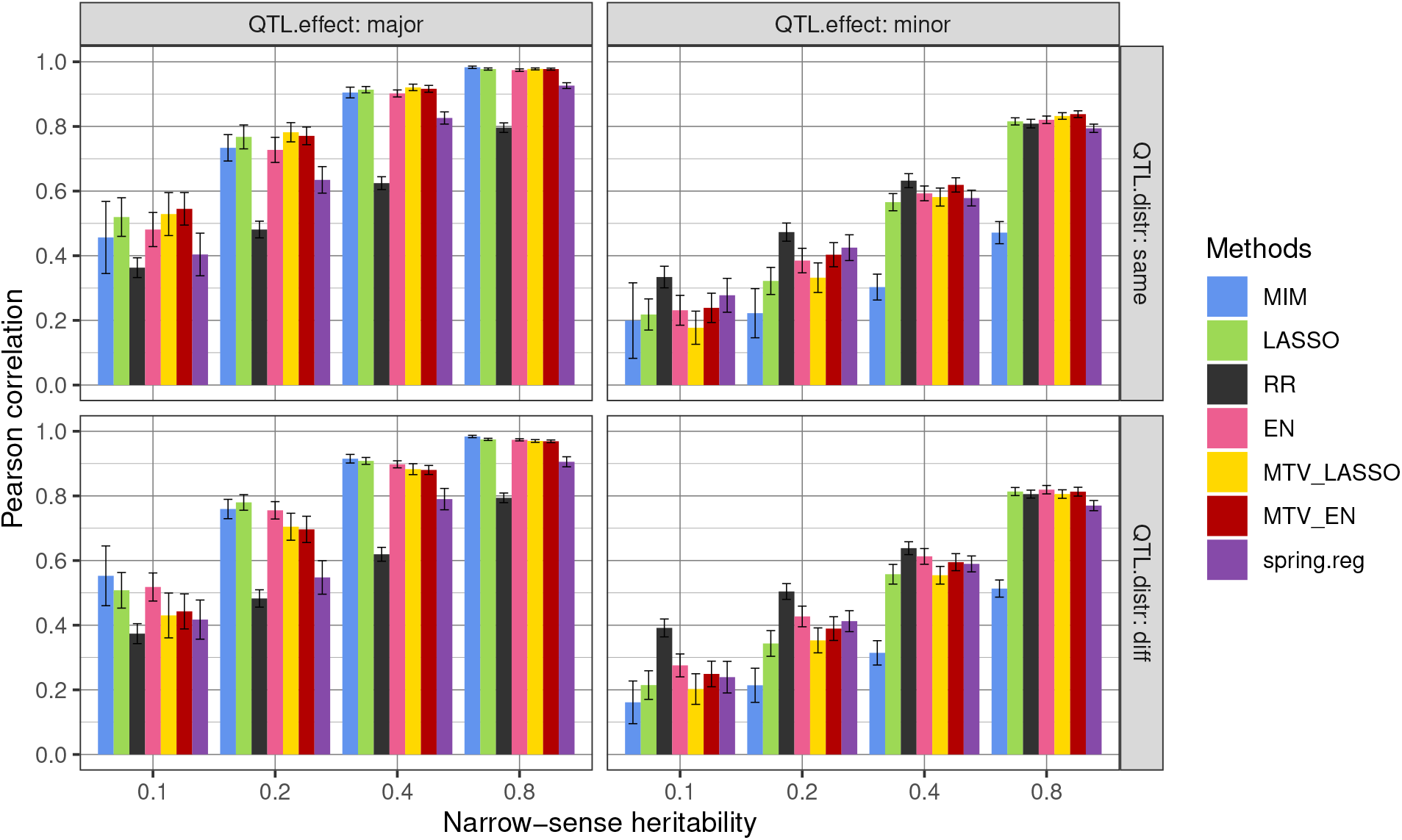
Genomic prediction accuracy (Pearson’s correlation between predicted and true genotypic values) of seven methods applied to 3,961 markers and two simulated traits in a bi-parental population with different heritability values and four QTL configurations (number x distribution among traits). major: 2 QTLs; minor: 50 QTLs; same: QTLs at the same positions for both traits; diff: QTLs at different positions between traits. For each heritability value and configuration, prediction accuracy was averaged over 100 values (2 traits x 10 simulation replicates x 5 cross-validation folds). The error bar corresponds to the 95% confidence interval around the mean.

The prediction accuracy of four additional methods is shown in Figures S6 and prediction accuracy values, as well as other performance metrics (see Materials and Methods) are in Table S7. All interval mapping methods yielded equivalent prediction accuracy. LASSO.GB did not improve performance compared to LASSO. MTV_RR showed equivalent performance as univariate RR. Prediction accuracy with spring.dir.ols was always lower than with spring.reg, and even very low for “minor” configurations. With 100 or 1000 simulated QTLs (under both QTL distributions) the ranking of methods based on prediction accuracy did not change compared to “minor” configurations (Figure S8).

##### Traits with different heritability values

To further compare prediction accuracy of univariate and multivariate methods, we simulated two correlated traits with different heritability values, 0.1 and 0.5. MTV_LASSO performed slightly better than univariate LASSO for the lowest heritability trait; however, differences were not significant (Figure S9). On the opposite, prediction accuracy was lower with MTV_LASSO than with univariate LASSO for the highest heritability trait, reaching quite low values with 200 simulated QTLs. The same trends were also visible for MTV_EN and EN. MTV_RR never improved prediction compared to RR and spring.reg never performed better than RR.

Since these results were unexpected, we also compared prediction accuracy of the above methods with the simulated data published by Jia and Jannink (2012). We obtained very similar differences among methods as with our simulated data, even though prediction accuracy was higher in all cases (Figure S10).

#### QTL detection: ROC curve results

We compared the main methods mentioned above (except RR which does not perform marker selection), as well as some robust extensions, for their marker selection performance with ROC curves, using the same simulated data (Figure 3) in the four configurations. On ROC curves, the closer a method gets to the optimum point (i.e. FPR =0 and TPR=1), the better. As expected, interval mapping methods (SIM and MIM) showed low selection performance when many minor QTLs were simulated and high selection performance when few major QTLs were simulated. Note that the MIM curve was hardly visible; it roughly overlapped with the SIM curve but stopped at a low FPR because it could not select many QTLs by design.

**Figure 3.**
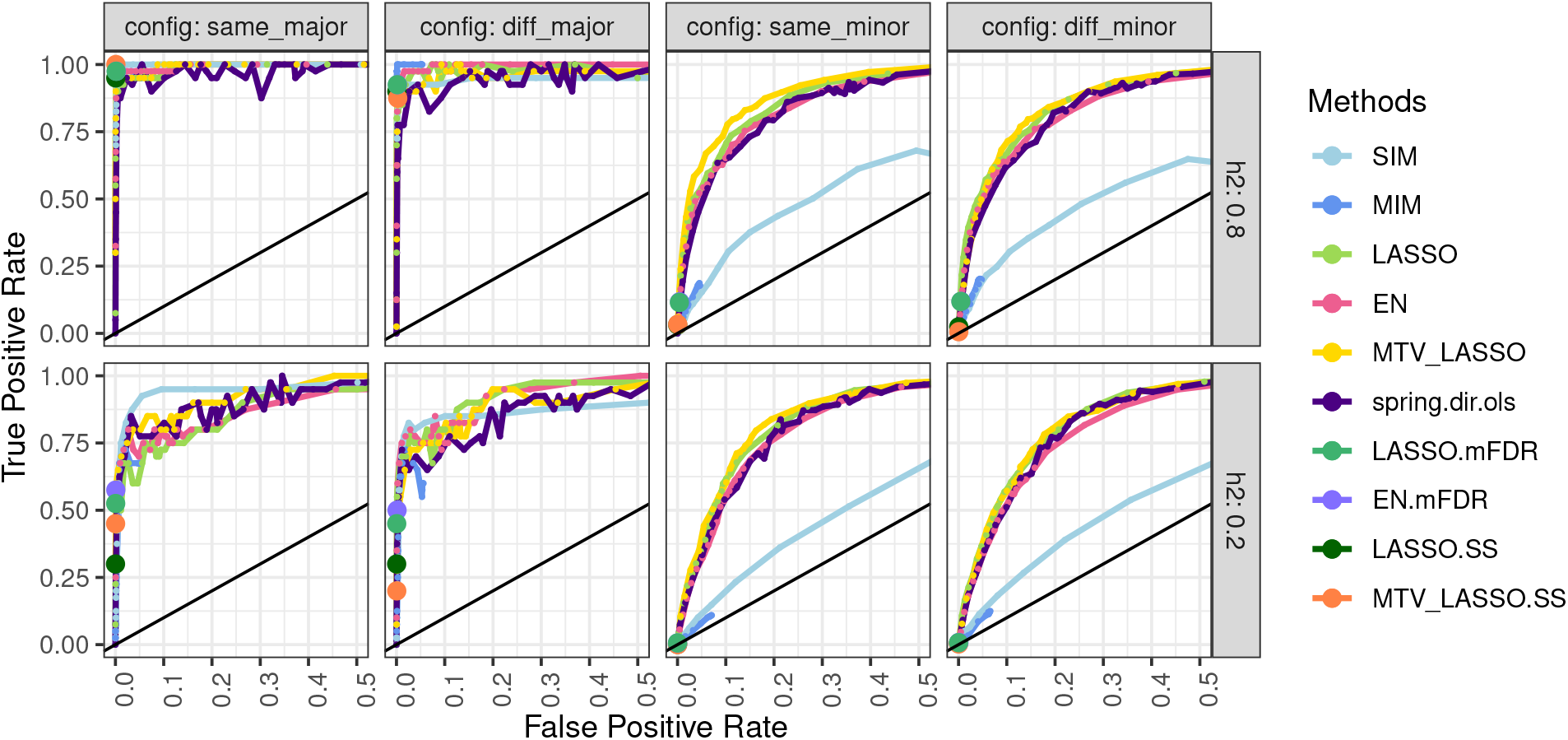
ROC curves for 10 methods applied to 3,961 markers and two simulated traits in a bi-parental population with two heritability values and four QTL configurations (number x distribution among traits). major: 2 QTLs; minor: 50 QTLs; same: QTLs at the same positions for both traits; diff: QTLs at different positions between traits. Results are averaged over 2 traits x 10 simulation replicates. TPR: True Positive Rate (number of correctly found QTLs / number of simulated QTLs), FPR: False Positive Rate (number of falsely found QTLs / number of markers outside a QTL). For robust methods (mFDR and SS), as the FPR remained very low, we display only a single point corresponding to the lowest parameter constraint and thus to the highest TPR.

The penalized regression methods always performed at least as well as the interval mapping methods or even much better in the case of “minor” configurations. Among penalized methods, no method was clearly better than the others in all configurations, except for a slight superiority of MTV_LASSO in the “same_minor” configuration. These methods, and particularly spring.dir.ols, displayed a high variability in classification results for two simulated QTLs (“major” configurations). Indeed, when one QTL was not detected among the two traits, there was a stronger impact on the TPR than with 50 simulated QTLs.

The most interesting part of the ROC curve for QTL detection is the left most part, i.e. with a low FPR (e.g. below 0.1). We hence calculated the partial Area Under the Curve (pAUC) for FPR between 0 and 0.1 for methods reaching that value (Figure S11). EN resulted in constantly high pAUC across configurations and heritability values. In contrast, pAUC for SIM was quite high at low heritability values for the “same_major” configuration but dropped for other configurations and heritability values.

### Results on experimental data

#### Computation of genotypic BLUPs

We first recomputed the genotypic BLUPs from the raw phenotypic data from (Coupel-Ledru *et al.* 2014, 2016) in order to control the model selection step in a reproducible way. These new BLUPs had a strong linear correlation (> 0.9) with those used in Coupel-Ledru *et al.* (2014, 2016), as shown in Figure S12. Note that in Coupel-Ledru *et al.* (2014), no BLUP was available for *DeltaPsi* and *PsiM* for WW condition because the genotype random effect was not selected (H²=0).

The estimates of broad-sense heritability followed the same trend as in Coupel-Ledru *et al.* (2014, 2016) (Figure S13). Nevertheless, values were not equal because we did not use exactly the same formula to estimate heritability. All the information about fitting linear mixed models (percentage of missing data, transformation applied if any, effects included in the selected model, residual variance, heritability estimate, coefficient of variation estimate and precision) is available in the first, online repository. Broad-sense heritability estimates were higher in WD condition than in WW for all traits except *DeltaBiomass*.

Genetic correlation between traits varied widely, some absolute correlation values being very high (e.g. up to 0.99 between *PsiM* and *DeltaPsi* in both conditions) because some traits derived from others (Figure S14).

#### Genomic predictive ability

Mean genomic predictive ability per trait ranged from −0.10 to 0.68 ((Figure 4 and Table S15). It decreased with broadsense heritability. IM methods (in blue) were always among the three worst methods for prediction. Based on the mean predictive ability averaged across traits, MTV_EN yielded the highest correlation (0.384), followed by RR (0.3721), MTV_RR (0.3716), MTV_LASSO (0.369), EN (0.357), spring.reg (0.344), LASSO (0.329), LASSO.GB (0.313), MIM (0.200) and SIM (0.162). However, based on the number of traits for which each method gave the best prediction, spring.reg had the highest score, with 6 traits out of 14, followed by MTV_EN (3 out of 14) and EN (2 out of 14).

**Figure 4.**
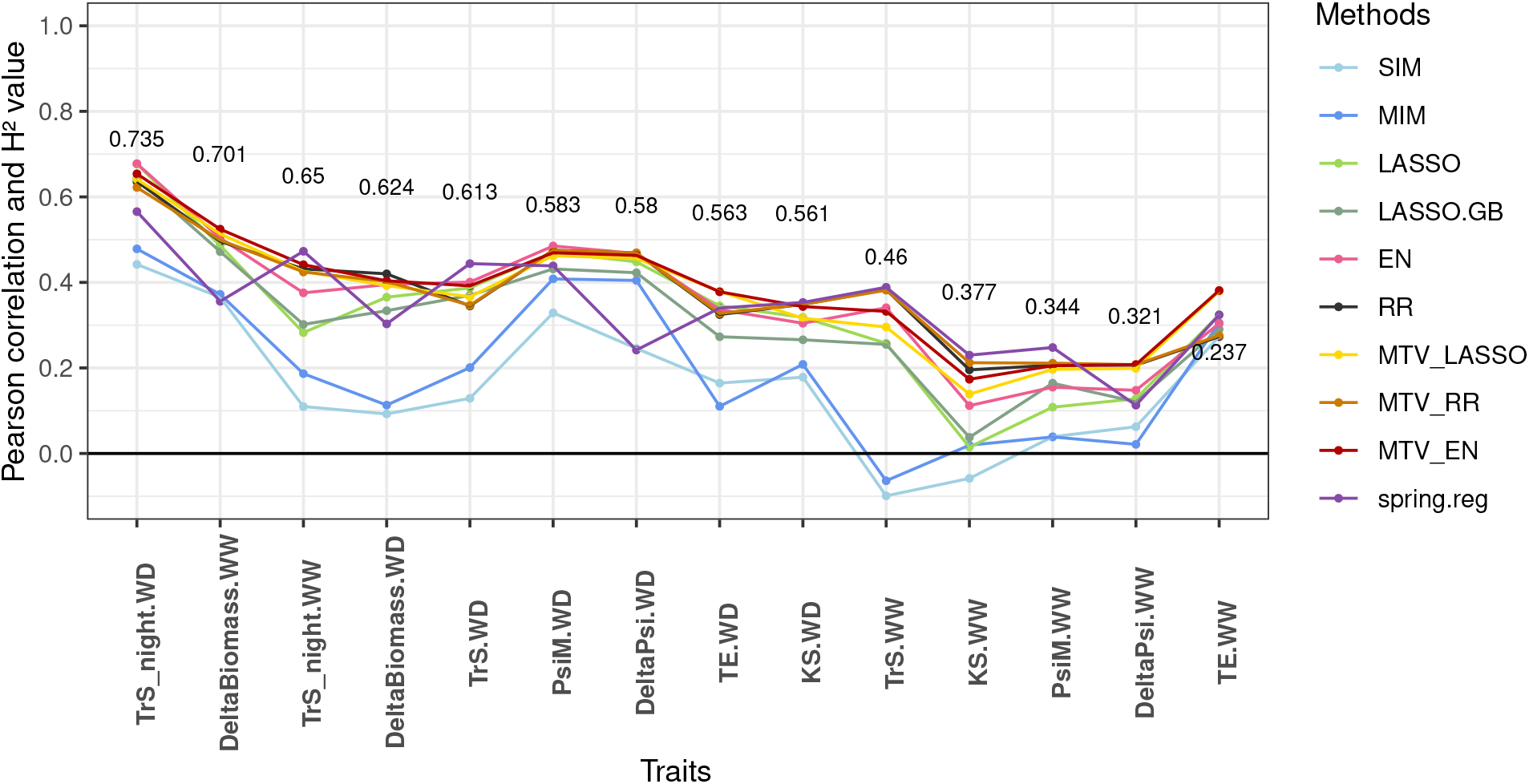
Mean genomic predictive ability (Pearson’s correlation between genotypic BLUPs and their predicted values), obtained by cross-validation for 10 methods applied to 14 traits related to water deficit and GBS gene-dose data, within a grapevine bi-parental population. Broad-sense heritability values are reported for each trait (y-position of the number corresponds to heritability estimate). Traits are ordered by decreasing heritability. For each trait, predictive ability is averaged over 10 cross-validation replicates x 5 cross-validation folds).

In a nutshell, MTV_EN and RR, tied with MTV_RR, provided the best mean predictive ability across traits. Even though spring.reg outperformed them for some traits, its performance was unstable, and especially low for *DeltaBiomass.WW*, *DeltaBiomass.WD*, *DeltaPsi.WW* and *DeltaPsi.WD*. For computational reasons, all traits could not be analyzed together with spring.reg, but were divided into three groups. These four traits with low predictive ability belonged to the same group. Yet, the effect of group membership on predictive ability was not significant at 5% (*p-value*=0.30 and percentage of variance explained=24%).

#### QTL detection

##### Interval mapping methods: comparison with previous results

For the 14 traits we analyzed, 26 QTLs were detected in Coupel-Ledru *et al.* (2014, 2016) using Composite Interval Mapping (CIM) on the SSR map. In comparison, using Multiple Interval Mapping (MIM), we found 21 QTLs on the SSR map and 25 with MIM on the SNP map (Figure S16).

Based on physical positions, we found 13 new QTLs (i.e. with non-overlapping CIM SSR QTLs physical positions) (Table S17) on six chromosomes for eight traits, and confirmed 21 of the 26 published QTLs, with a notable reduction of QTL intervals on chromosome 13 (Figure S16). The 15 QTLs found by all three methods (CIM SSR, MIM SSR and MIM SNP) explained the highest mean percentage of variance (Figure S18).

##### Comparison of marker selection among a subset of methods

After applying 11 methods for SNP selection (Table S19), we performed a first comparison of marker selection between MIM, as the reference method for QTL detection, and both LASSO and EN, because our simulation results showed that they selected relevant markers in various genetic determinism configurations (Figures 3 and S11).

The number of markers selected by MIM, LASSO and EN was 905, 1009 and 1550, respectively (Table S19). For each trait, MIM identified markers on a small number of chromosomes (from 0 to 5), while both EN and LASSO selected markers on many chromosomes (from 6 to 19, Table S19). The number of selected markers per trait seemed partly linked to trait heritability: more markers were selected when heritability was high (Figures 4 and 5). More markers were selected by EN than by other methods for all traits (except for *KS.WW*). Nearly all markers selected by LASSO were also selected by EN (954 out of 1009), i.e., there were only few markers selected by LASSO only. MIM selection was quite different from LASSO and EN selections (184 out of 905 were common with EN, LASSO or both) but most markers selected by MIM and at least one penalized method were selected by both EN and LASSO. The number of markers selected by EN and MIM ranged from 0 to 59 over traits, with a median value of 16.

**Figure 5.**
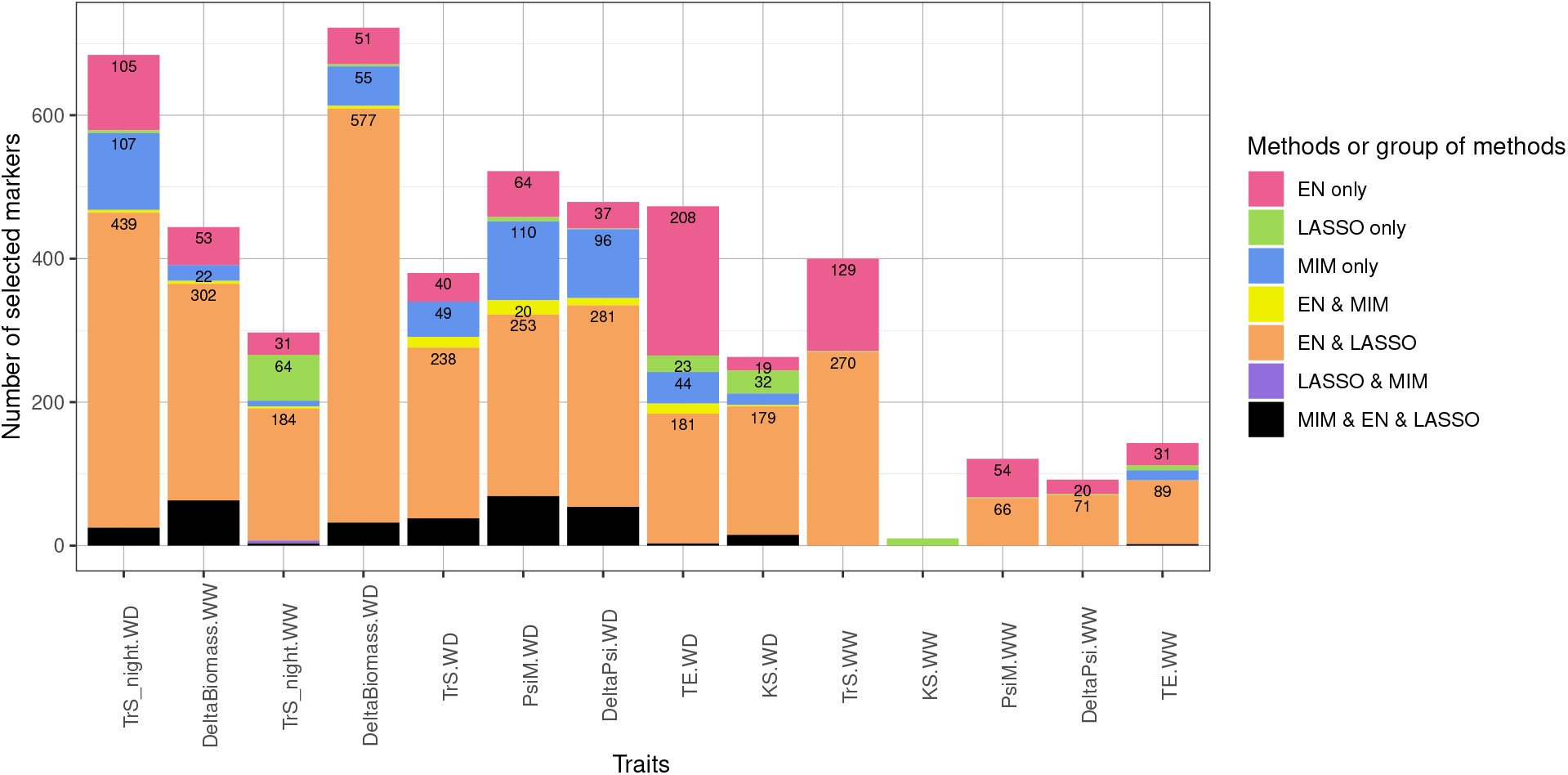
Number of selected markers per method or group of methods, for three methods applied to 14 traits related to water deficit and GBS gene-dose data, within a grapevine bi-parental population. Traits are ordered as in Figure 4. Number of selected markers (extended to markers in high LD, see Materials and Methods) per category are indicated at the top of each rectangle. Methods followed by « only » are for the number of markers selected by this method that are not selected by any of the two other methods (among EN, LASSO and MIM).

##### Determination of highly reliable QTLs

To address the intersection of SNP selection by all methods, and determine the number of reliable intervals (QTLs) and their position, we examined in more detail marker selection for each trait and chromosome. Detailed results, including genetic and physical positions and the percentage of variance explained, are given in Table S19. A visualization of these results is given in Figure 6 for night-time transpiration under water deficit (*TrS_night.WD*) and in Figure S20 for all traits.

**Figure 6.**
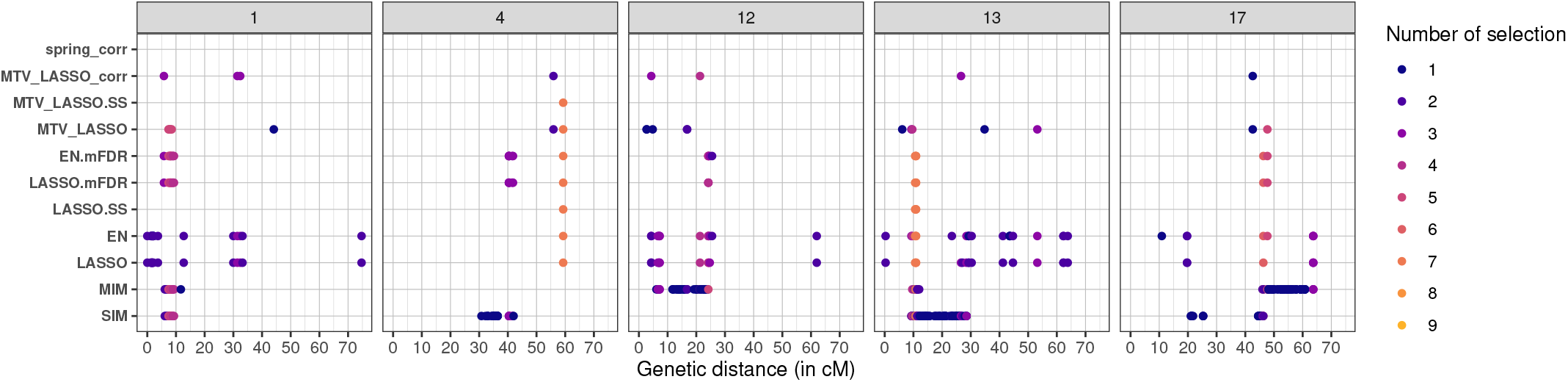
Marker selection by all methods for *TrS_night.WD* trait on chromosomes 1,4,12,13 and 17. Each marker selected by a given method is represented by a colored point, the color indicating the number of methods that have selected that specific marker. The boxes correspond to chromosomes and the x-axis to the position along the genetic map (in cM).

**Figure 7.**
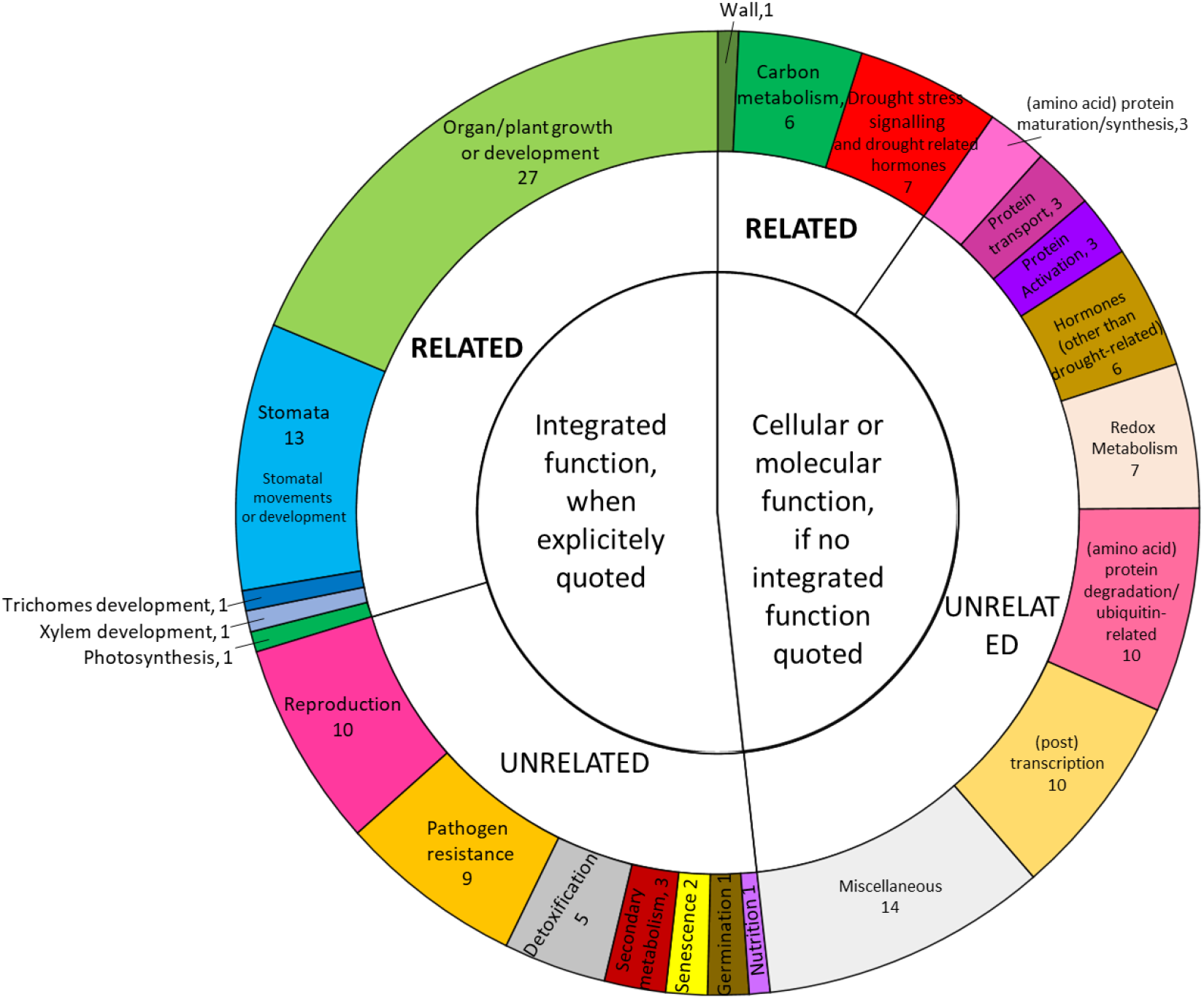
Functional classification of the annotated genes underlying the highly reliable QTL detected on chromosome 4 for night-time transpiration, growth and transpiration efficiency. Hierarchical classification of the 161 genes based on their functions. See Table S25 for the details of this classification. When an integrated function at the organ or plant level was explicitly quoted in the gene annotation, genes were classified on this basis. When no integrated function was explicitly quoted, they were classified based on their cellular or molecular function. In both cases, functions were then classified as “Related” if related to the traits of interest in this QTL, or “Unrelated” if not.

Most of the time, more markers were selected for traits under water deficit than for traits in well-watered conditions, and they were most often selected by several methods. We showed that penalized methods tend to select the same markers, not only close ones; for example, for *TrS*_*night*.*WD* on chromosome 4, the same marker (at physical position 21,079,664 bp) was selected by seven methods (Figure 6).

We considered markers selected by both MIM and EN.mFDR as highly reliable ones for three reasons: 1) markers selected by both MIM and EN were considered as reliable ones (see above); 2) simulations showed that MIM and mFDR methods led to a very low FPR; 3) these methods belong to different method classes (interval mapping *vs* penalized regression). We also considered as highly reliable the markers selected by at least five methods. These criteria resulted in a set of 59 highly reliable selected markers, which were converted to genetic intervals of ± 3 cM around each selected marker. Overlapping intervals per trait were merged, resulting in 25 highly reliable QTLs.

These 25 QTLs involved nine traits, mostly under water deficit, and were located on seven chromosomes (Figure S21). QTLs colocalized for different traits, such as on chromosome 1, had similar distributions of genotypic BLUPs according to genotypic classes (Figure S22).

Among these 25 QTLs, 16 had overlapping physical intervals with CIM SSR QTLs and one was very close to a CIM SSR QTL (details about these 25 QTLs are in Table S23). Thus, we found eight new highly reliable QTLs, among which five were not detected by MIM. In particular, a completely new QTL for *TrS_night.WD* was found alone on chromosome 12. Most other new QTLs were colocalized with previously found QTLs in single year analysis and/or for the other watering condition. Notably, we observed colocalization of *TrS_night.WD*, *TE.WD* and *DeltaBiomass.WD* QTLs on chromosomes 4 and 17.

In total, the percentage of variance explained (adjusted R²) per trait was 51.28% for *TrS_night.WD* (36% in 2012 for Coupel-Ledru *et al.* (2016), 33.88% for *PsiM.WD*, 31.41% for *DeltaPsi.WD*, 26.88% for *DeltaBiomass.WW*, 19.38% for *TE.WD*, 18.62% for *TE.WW*, 16.99% for *KS.WD*, 14.88% for *DeltaBiomass.WD* and 8.55% for *TrS.WD*.

#### Candidate genes

After merging the QTLs colocalized between traits, we obtained 12 intervals, located on chromosomes 1, 4, 10, 12, 13, 17 and 18, harboring a total of 3,461 genes according to the VCost.v3 annotation (Canaguier *et al.* 2017). Among them, 2,379 had a NCBI Refseq identifier and 1,757 a putative function (Table S24). We then focused our analysis on the eight “new” intervals, i.e. those which were not overlapping with CIM SSR intervals. They encompassed 1,155 genes, half of which were annotated. We were able to retrieve from TAIR and/or UniProt a more precise description of the genes function for 86% of the annotated genes (Table S24). The remaining ones either did not have any homologous gene in Arabidopsis thaliana or were not described in the above-mentioned databases. RNA-seq data was available on the GREAT platform for 90% of the annotated genes. We further focused our analysis on the highly reliable QTL co-located on chromosome 4 for *TE*, *TrS_night* and *DeltaBiomass* under various conditions. We proceeded to a functional classification of the 161 annotated genes underlying this QTL, based on the full description previously retrieved (Table S25 and 7). For 75 genes, an integrated function at the plant or organ level was explicitly quoted in the description. We grouped these integrated functions into 12 major groups: stomata, trichomes development, xylem development, growth or development, photosynthesis, wall, reproduction, pathogen resistance, detoxification, secondary metabolism, senescence, germination, and nutrition. A substantial number of genes were related to the functions of major interest in relation to the traits for which QTLs co-localized on this chromosome: 15 genes related to hydraulics (stomata, xylem, trichomes), relevant for *TrS_night* and thus *TE*; 27 to growth or development and one to photosynthesis, both relevant to *DeltaBiomass* and thus *TE*. For the 86 genes for which an integrated function was not explicitly quoted, we further built a classification based on their cellular or molecular function. Among them, we found six genes related to carbon metabolism, one to wall formation (both relevant for *DeltaBiomass*) and six to drought stress signaling and drought related hormones (relevant for *TrS_night*).

## Discussion

To provide new insights into the complex genetic determinism of vegetative traits under different watering conditions, the contributions of this study are three-fold. We compared by simulation several univariate and multivariate methods for genomic prediction and QTL detection, increased the density of genotyping data, and re-analyzed grapevine phenotypes obtained under semi-controlled conditions. In particular, we showed that penalized methods are valuable not only for prediction but also for QTL detection. Indeed, we found new QTL using these methods and identified relevant candidate genes.

### Methodological aspects: method comparisons

#### Handling linkage disequilibrium

Interval mapping methods estimate genotypic probabilities between markers according to a genetic map which is computationally costly to build. On the other hand, most penalized methods do not require any previous knowledge on LD.

The LASSO assumption that all predictor variables are independent is all the more violated that there are many markers. In the case of a group of correlated predictors (e.g., SNPs in LD), EN selects either none or all predictors within the group with close estimated values (Zou and Hastie 2005) whereas LASSO selects a single predictor. In that sense, EN aims at correcting the drawbacks of LASSO when predictor variables are highly correlated. By exploring a large number of configurations of the finite-sample high-dimensional regression problem, Wang *et al.* (2020) showed that EN is competitive for both prediction and selection in most cases with highly correlated predictors. In agreement with these results, we showed that EN performed well for both prediction and selection on our simulated data, and that multivariate EN performed the best for prediction on the grapevine experimental data.

We also compared SPRING that can explicitly make use of a genetic map. We observed that SPRING had a larger increase in predictive ability from SSR to SNP design matrix than other methods (Figure S4). This was probably due to the fact that SPRING uses LD pattern for prediction, this pattern being better captured with a dense genetic map. However, SPRING showed no systematic advantage over other penalized methods for prediction with the dense SNP map (Figures 2, 4).

#### Comparison between interval-mapping and penalized regression methods for genomic prediction

As expected, IM methods performed poorly to predict accurate genotypic values when QTL number was large (Bernardo and Yu 2007; Lorenzana and Bernardo 2009; Mayor and Bernardo 2009; Olatoye *et al.* 2019) (Figures 2 and S6). Therefore, for complex traits, genomic prediction should not be based only on QTLs detected by IM methods.

Among univariate penalized methods, none performed best in all cases (Figures 2, 4 and S6), as also found in the literature (Riedelsheimer *et al.* 2012; Heslot *et al.* 2012; Azodi *et al.* 2019). As shown by simulation, RR was better adapted to highly polygenic genetic architecture whereas LASSO was better adapted to a few major QTLs. Moreover, in the case of many minor QTLs, RR was the most stable method across heritability values, as previously described for several traits and species (Heslot *et al.* 2012; Azodi *et al.* 2019). However, RR prediction accuracy dropped when QTL number was too small whereas EN still predicted as well as LASSO. EN was hence well adapted to various numbers and distributions of QTLs.

##### Multivariate *vs* univariate

When the same heritability was simulated for both trait variables, no superiority of multivariate methods was observed, even when both traits had QTLs at the same positions (Figures 2 and S6).When different heritability values were simulated for the two traits, we observed a slight superiority of MTV_LASSO (resp. MTV_EN) over LASSO (resp. EN) only in the “same” and “major” configuration (with both traits sharing the same two QTLs) for the trait with small heritability (Figure S9).

Other authors which tested multivariate GP on simulated data systematically applied different heritability values and they found a superiority of multivariate methods over univariate ones for the trait with the smallest heritability (Calus and Veerkamp 2011; Guo *et al.* 2014; Jiang *et al.* 2015; Dagnachew and Meuwissen 2019). However, all these studies were based on a smaller, more favorable, *p*/*n* ratio, a key component of high-dimensional models (Verzelen 2012). For example, in Jia and Jannink (2012), their 500 observations for 2,020 predictors correspond to a ratio of ~ 4, compared to our 188 observations for 3,961 predictors corresponding to a ratio of ~ 21. Indeed, parameters *n* and *p* are involved in the sample complexity function defined in Obozinski *et al.* (2011), which predicts the theoretical cases where the MTV_LASSO is superior to its univariate counterpart in terms of variable selection. Accordingly, applying our methods on Jia and Jannink (2012) data allowed us to display a higher difference between univariate and multivariate LASSO than with our simulated data.

Unexpectedly, when reanalyzing the data simulated by Jia and Jannink (2012), we obtained lower prediction accuracy with our MTV_LASSO (Figure S11) than they did with their multivariate BayesA (their Figure 1A). A similar result in a univariate setting was found by Guan and Stephens (2011) who compared BSVR (comparable to BayesA) and the LASSO. They found that BSVR had a markedly higher power than the LASSO. Moreover, the parameters of both BSVR (in Guan and Stephens (2011)) and BayesA (in Jia and Jannink (2012)) were estimated with a MCMC algorithm. No inner cross-validation was needed, hence the sample used to train the model was larger. This difference may explain why Figure 1A from Jia and Jannink (2012) shows better prediction accuracies for multi-trait models compared to their single-trait counterparts, although their figure did not display any confidence interval. Note that our RR prediction accuracies were close to those of their GBLUP (univariate and multivariate). As a conclusion, prediction accuracy is affected both by the dimension of the problem (i.e., *n* and *p*) and the method used (i.e., Bayesian with MCMC or cross-validation).

For experimental data, we observed that MTV_LASSO (respectively MTV_EN) was superior to LASSO (resp. EN) for the five traits with the smallest heritability (Figure 4). This improvement suggests that MTV_LASSO (resp. MTV_EN) was able to borrow signal from the most heritable traits to the least heritable ones, likely because of a genetic architecture partially overlapping between these traits. This interpretation is reinforced by the fact that a QTL for low-H2 trait, *TE.WW*, colocalizes on chromosome 4 with QTLs for four high-to-moderate-H2 traits (*TrS_night.WD*, *DeltaBiomass.WW*, *DeltaBiomass.WD* and *TE.WD*). This improvement was not found in Jia and Jannink (2012), who also tested their methods on real pine data from Resende *et al.* (2012). These observations suggest that the number of traits analyzed (14 in our case and 2 in Jia and Jannink (2012) study) may also play a role in the prediction accuracy of multivariate over univariate methods.

#### Comparison between interval-mapping and penalized regression methods for QTL detection

To the best of our knowledge, comparison with the ROC curve between IM and penalized regression methods has never been done before in terms of marker selection. Other publications (Cho *et al.* 2010; Li and Sillanpää 2012; Waldmann *et al.* 2013) successfully applied LASSO or EN for performing GWAS, but none of them compared IM and penalized methods for QTL identification. As expected, we found that IM methods are adapted to detect a few major QTLs but not many minor QTLs (Figure 3). Moreover, we found that penalized methods could be as good at marker selection as IM methods, and even far better when there are many minor QTLs. Among the penalized methods we compared, none clearly outperformed the others for marker selection in all configurations.

##### Multivariate *vs* univariate

As the MTV_LASSO selects one predictor for all traits, its superiority over univariate LASSO depends on QTL distribution across traits, notably on the amount of genetic basis shared by the traits (Obozinski *et al.* 2011). However, as for prediction, we showed that MTV_LASSO performance was not different whether QTLs were at the same or at different positions across traits (Figure 3). Nevertheless, we observed that MTV_LASSO was slightly better than LASSO when many QTLs were simulated. SPRING had never been evaluated before for its quality of predictor selection. As for prediction, SPRING showed unstable results across our simulation replicates and hyper-parameter values. However, for the ROC curve, we did not include predictor structure in the model, which may hamper marker selection quality.

#### Efficient default method for both QTL detection and genomic prediction

IM methods were designed for marker selection; hence they are not expected to be optimal for prediction, and we confirmed that. Among penalized regression methods, some may be better at prediction than marker selection, and vice versa. For example, our results showed that EN performed well across several configurations for both aims. Some methods such as SPRING are specially adapted to handle both purposes but it gave too variable results for either prediction or QTL detection. However, SPRING is a recent method that still can be improved in order to correct this drawback.

New penalized regression methods are continuously being developed. In particular, graph structured sparse subset selection (Grass) recently proved to outperform existing methods for both prediction and predictor selection, thanks to a *L*0 regularization that limits the number of nonzero coefficients in the model (Do *et al.* 2020). It could be tested on our data when its implementation becomes available. Moreover, multivariate methods are presented as being more efficient at using the whole signal in the data, whether for marker selection (Inouye *et al.* 2012) or prediction (Jia and Jannink 2012; Guo *et al.* 2014), but our results revealed no systematic advantage of multivariate methods over univariate ones for both aims.

Using penalized methods for both marker selection and genomic prediction requires adapted hyper-parameter values. For EN, LASSO and SPRING, the *λ* value controls sparsity (e.g., the number of selected markers). Thus, the optimal value of *λ* might not be the same if the aim is to limit the FPR or to maximize the predictive ability (Li and Sillanpää 2012). For prediction, we traditionally use cross-validation to tune hyper-parameters by minimizing MSE. For marker selection, there is no direct equivalence. That is why we tested extensions of these methods (mFDR and SS) which control sparsity for robust marker selection and they proved to be efficient to select the most relevant markers.

In order to shed light on the link between prediction accuracy and marker selection, we plotted the prediction accuracy at each point of the ROC curve for EN and EN.mFDR against FPR for minor configurations (with 50 simulated QTLs) (Figure S26). For EN, we showed that prediction accuracy reached its maximum when FPR was below 0.05. Then, FPR increased while prediction accuracy decreased, until it reached a plateau. This means that prediction quality is intimately linked to selection quality, especially at low heritability. For EN.mFDR, the FPR stayed always below 0.015 but the prediction accuracy was lower.

As a consequence, as an efficient default method, we advise at this stage to apply EN for performing genomic prediction, and its extension EN.mFDR for performing sparser marker selection.

#### Genetic determinism and prediction of grapevine response to water deficit

Based on experimental data on the Syrah x Grenache progeny (new genotypic data and already published phenotypic data), we compared the same methods as above for both prediction and marker selection. To the best of our knowledge, grapevine GP within a bi-parental family has been applied only to a limited number of traits, with very few methods and never using multivariate GP. Fodor *et al.* (2014) studied GP in grapevine with simulated data on a diverse and structured population, they tested RR-BLUP, Bayesian Lasso, and a combination of marker selection and RR. Viana *et al.* (2016a) used an inter-specific grapevine bi-parental population. They predicted cluster and berry phenotypes (number and length of clusters, number of berries, berry weight, juice pH, titrable acidity) with RR-BLUP and Bayesian LASSO applied to table grape breeding. In addition to yielding further insights into method comparison beyond those obtained by simulation, our study brought valuable novel biological knowledge about grapevine water use under different watering conditions. Indeed, new methods and the new SNP genetic map allowed us to find novel QTLs, as compared to those previously detected with the same phenotypic data (Coupel-Ledru *et al.* 2014, 2016).

#### Predictive ability and genetic architecture

Among univariate penalized methods, RR generally had equivalent or better predictive ability than LASSO. For the traits with the largest discrepancy between RR and LASSO, this suggests that trait variability was rather due to many minor QTLs rather than to a few major ones. On the other hand, predictive abilities of sparse methods (e.g. LASSO and IM methods) were better than RR for *PsiM.WD*, *DeltaPsi.WD* and *TE.WW* traits, suggesting a more major genetic architecture. We observed that some genomic regions were less densely covered by the SNP genetic map (e.g., a 10 cM gap on chromosome 19), which might lead to a decrease in predictive ability for traits with QTLs in these regions. We tested this hypothesis for penalized methods, by using the raw genotypic data imputed with the mean (SNP.raw on Figure S4). For most traits, this design matrix gave worse predictions than with other SNP ones, except for *TE.WW*, for which the raw matrix gave the best predictive abilities (data not shown). This suggests that some QTLs for *TE.WW* were lost (markers not selected) when we predicted with sparser design matrices, whereas this was not the case for other traits. Filtering markers by genetic mapping for prediction purpose thus proved to be useful for most traits.

Furthermore, we tested several design matrices for GP on experimental data. The matrices derived from the SNP map led to better predictive ability than those derived from the SSR map, due to higher density, while the additive + dominant coding of allelic effects did not provide any increase in predictive ability (Figure S4). This could suggest that dominance effects have negligible impact on these traits. Nevertheless, the additive + dominant coding double the matrix dimension (up to 31,688 predictors), which might hamper allelic effect estimation and thus, prediction. Finally, non-additive genetic effects such as epistasis could be involved while not considered by the penalized methods used. We therefore tested the superiority of LASSO.GB over LASSO. Extreme Gradient Boosting methods are indeed among the best machine learning methods (Chen and Guestrin 2016). LASSO.GB did not markedly increase predictive ability on experimental data (Figure 4). However, we cannot exclude that this might be due to a poor optimization of Extreme Gradient Boosting parameters or to insufficient number of observations to correctly fit the model.

#### Candidate gene analysis

The thorough methodology deployed for candidate genes analysis allowed us not only to retrieve a list of the genes underlying the QTLs of interest, but also to classify them based on their function and expression in order to point at more likely candidates. We focused on the highly reliable QTL detected on chromosome 4 for *TrS_night*, *TE* and *DeltaBiomass*. *TrS_night* QTL was previously described as a promising target for marker assisted selection, as alleles limiting night-time transpiration also favor plant growth, resulting in a double, beneficial impact on improving transpiration efficiency (Coupel-Ledru *et al.* 2016). Moreover, this QTL was found by seven methods. Within a plethora of integrated functions represented within the list of annotated genes underlying this QTL, we show here that a subset of more likely candidates can be defined as possibly related to the traits of interest. These include on one hand, genes related to broad-sense hydraulics and water loss, with a possible direct impact on *TrS_night*: seven genes involved in stomatal development, nine genes involved in stomatal opening -sometimes through the abscicic acid signalling pathway-, one to xylem development and one to trichome development (Table S25). One of these genes, the trihelix transcription factor GT-2 (Vitvi04g01604), was specifically shown to impact transpiration and transpiration efficiency in Arabidopsis by acting as a negative regulator of stomatal density. On the other hand, 27 genes among the list are directly related to growth, development, or photosynthesis, meaning a possible direct impact on *DeltaBiomass*. A histidine kinase 1 (Vitvi04g01483) may be a particularly interesting candidate for its multiple roles in ABA signalling, stomatal development and plant growth known in Arabidopsis, hence potentially simultaneously acting on both components of *TE*. Both these likely candidates were often highly expressed in grapevine leaves according to the data retrieved from the RNA-seq database. The reduction of confidence interval did drastically reduce the number of genes as well as the subsequent analyses, but the list is still extensive. More precise analyses of these candidate genes, including functional genomic work and possible gene editing of some of them will be now necessary to identify the genes under these new QTLs.

## Conclusion

Faced with the threat of climate change and the challenge of decreasing inputs while maintaining yield and quality, deciphering the genetic architecture of target traits is a most needed endeavor. In this goal of importance to all agricultural species whatever the traits under investigation, the approach developed in this article aimed at harnessing the most information as possible from dense genotyping and accurate phenotypic data. Among the wealth of available methods, we focused our comparison on univariate *vs* multivariate ones. Moreover, rather than decoupling genomic prediction from the identification of major QTLs, we argue for the need to purse both goals jointly. Indeed, they provide complementary information on the genetic architecture of the target traits as well the key functions underlying them. As such, we provided an in-depth investigation mobilizing both simulated and experimental data, hence of interest beyond our grapevine case study, hoping that it will contribute to a way forward to other researchers working on other species. Of interest to quantitative geneticists, our results notably emphasized the interest of the Elastic Net, available as both a univariate and a multivariate version, as an efficient, default method for genomic prediction, followed by the mFDR control for the robust identification of QTLs. Moreover, of interest to plant biologists who seek to understand the response to water stress, our results highlighted several candidate genes underlying the integrated traits of night-time transpiration, transpiration efficiency and biomass production. For some of them, their functions confirm and suggest causal links with stomatal functioning, trichome development or the ABA pathway.

## Acknowledgments

We thank Marie Perrot-Dockès for her help concerning penalized regression methods, the South Green platform for computing, notably Bertrand Pitollat, and the GREAT platform, notably Amandine Velt and Camille Rustenholz. We also acknowledge the funding of the SelVi and FruitSelGen projects (BAP department and Selgen metaprogram of INRAE). Partial funding of the PhD was provided by ANRT (grant number 2018/0577), IFV and Inter-Rhône.

